# Molecular Mechanism for Rotational Switching of the Bacterial Flagellar Motor

**DOI:** 10.1101/2020.05.18.101634

**Authors:** Yunjie Chang, Kai Zhang, Brittany L. Carroll, Xiaowei Zhao, Nyles W. Charon, Steven J. Norris, Md A Motaleb, Chunhao Li, Jun Liu

## Abstract

The bacterial flagellar motor is a remarkable nanomachine that can rapidly rotate in both counter-clockwise (CCW) and clockwise (CW) senses. The transitions between CCW and CW rotation are critical for chemotaxis, and they are controlled by a signaling protein (CheY-P) that interacts with a switch complex at the cytoplasmic side of the flagellar motor. However, the exact molecular mechanism by which CheY-P controls the motor rotational switch remains enigmatic. Here, we use the Lyme disease spirochete, *Borrelia burgdorferi*, as the model system to dissect the mechanism underlying flagellar rotational switching. We first determined high resolution *in situ* motor structures in the *cheX* and *cheY3* mutants in which motors are genetically locked in CCW or CW rotation. The structures showed that the CheY3 protein of *B. burgdorferi* interacts directly with the FliM protein of the switch complex in a phosphorylation-dependent manner. The binding of CheY3-P to FliM induces a major remodeling of the switch protein FliG2 that alters its interaction with the torque generator. Because the remodeling of FliG2 is directly correlated with the rotational direction, our data lead to a model for flagellar function in which the torque generator rotates in response to an inward flow of H^+^ driven by the proton motive force. Rapid conformational changes of FliG2 allow the switch complex to interact with opposite sides of the rotating torque generator, thereby facilitating rotational switching between CW and CCW.

## Introduction

The bacterial flagellum is a remarkable nanomachine that can rotate in both the counter-clockwise (CCW) and clockwise (CW) directions and can switch rapidly between the two rotational states^1-4^. Regulation of the rotational direction is key for bacterial chemotaxis, a behavior that enables the cells to move toward attractants or away from repellents^5,6^. In externally flagellated bacteria, such as *Escherichia coli* and *Salmonella enterica* serovar Typhimurium, CCW rotation of the flagella coalesces the external helical flagellar filaments into a bundle that produces smooth swimming (a run), and CW rotation disrupts the bundle and reorients the cell (a tumble)(Extended Data Fig. 1**a, b**)^1^. A sophisticated chemotaxis signaling system allows the cell to sense chemical stimuli and transmit this information via a phosphorylated form of the response regulator CheY to regulate the direction of rotation^5,6^. Although it is well known that lower levels of CheY-P promote CCW rotation and higher levels promote CW rotation, the exact mechanism of CheY-P induced rotational switching is unknown^1,3,4^. Intriguingly, recent data suggest that flagellar switch proteins are highly dynamic and that the number of subunits vary significantly in *E. coli* and *S. enterica* motors rotating CCW and CW^7,8^. However, it is unclear how the flagella could accommodate such large changes while still maintaining rapid rotation and switching.

Spirochetes are a unique group of bacteria with distinct morphology and motility^9,10^. Spirochetes possess multiple internal periplasmic flagella (PF) that are attached near each cell pole. These flagella are located between the outer membrane sheath and the cell cylinder, and their rotation causes the entire cell body to rotate (Extended Data Fig. 1**e, f**)^9^. Spirochetes run when the anterior flagella rotate CCW and the posterior flagella rotate CW^11,12^. When the flagella at both poles rotate in the same direction, the spirochetes flex in place and fail to move translationally^11,12^. To swim toward an attractant, spirochetes have evolved a complex chemotaxis and motility system to coordinate rotation of the PF at the two cell poles^9,13^. CheY3 is a key response regulator that is essential for chemotaxis in *B. burgdorferi;* Δ*cheY3* mutant cells are non-chemotactic and constantly run^12^. CheX is the CheY-P phosphatase identified in *B. burgdorferi^14^*. A Δ*cheX* mutant constantly flexes and is not able to run or reverse *in vitro^14^.* How the CheY3-P coordinates flagellar rotation at both poles to achieve directional migration in chemical gradients is a key question in spirochete motility and chemotaxis^9,10^.

The rotary motor is the most intricate part of the flagellum; it is responsible for flagellar assembly, rotation and directional switching. Whereas details of the motor structures vary among species, the core components, which are the products of billions of years of evolution, are highly conserved^15,16^. The membrane-bound stator and the switch complex (also called C-ring) are directly responsible for flagellar rotation and switching. The stator complex is the torque generator powered by ion flux across the membrane. It is composed of two transmembrane proteins, called MotA and MotB in *E. coli* and *B. burgdorferi*^17,18^. MotA has a large cytoplasmic domain, which contains several conserved charged residues that are critical for the interaction with the switch complex^19^. MotB has a large periplasmic domain that is believed to bind to the peptidoglycan layer^20,21^. The switch complex is comprised of three proteins (FliG, FliM, and FliN) that assemble to form the characteristic C-ring at the cytoplasmic side of the motor. FliG is the protein most directly involved in interacting with the stator to generate torque^22^. In *B. burgdorferi*, which has two FliG proteins, FliG2 is present in the C-ring and plays a similar role as its counterpart in other bacteria. FliG1 is located at one cell pole; it remains unknown if it is a part of the C-ring^23^. FliM and FliN are extensively involved in switching the direction of the motor^3^.

Here, we deployed cryo-electron tomography (cryoET) to visualize the *B. burgdorferi* motors in Δ*cheX* and Δ*cheY3* mutants in which the flagella are locked in CCW and/or CW rotation. The resulting *in situ* structures of the stator complex and switch complex enable us to uncover that binding of CheY3-P to FliM induces a profound conformational change in FliG2 between CW and CCW rotation. Importantly, our data suggest a model in which the stator complexes rotate in response to proton flow and interact with FliG2 that are in radically different conformations to drive CW and CCW rotation.

## Results

### *In situ* structure of the flagellar motor in constantly flexing Δ*cheX* cells

Recent *in situ* structural analysis of the wild-type (WT) and Δ*motB* flagellar motors in *B. burgdorferi* demonstrates the utility of combining cryoET and genetic approaches for understanding the structure and function of the intact *B. burgdorferi* flagellar motor^18^. In an unsynchronized pool of WT cells, the motors constantly change their rotational senses to drive the spirochetal motility. Therefore, it is challenging to sort out the WT motors into distinct CW or CCW conformations. To overcome this problem, we analyzed the flagellar motors in Δ*cheX* mutant cells which continuously flex and unable to run or reverse *in vitro^14^* (Supplementary Video 1). Due to high levels of CheY3-P in the Δ*cheX* cells^14^, the motors in both cell tips are expected to be locked in CW rotation. Another advantage of analyzing the motors of these cells is that due to high levels of CheY3-P it may be possible to visualize the switch complex when it is occupied by this signaling protein.

To determine the *in situ* flagellar motor structure by cryoET and subtomogram averaging, we analyzed 1,056 flagellar motors from 246 tomograms of Δ*cheX* cell poles (Extended Data Fig. 2**a**, Extended Data Table 1). The averaged structure reveals the core components of the flagellar motor – such as the stator, C-ring, export apparatus, and spirochete-specific collar^18^ (Fig. 1**a**). A *B. burgdorferi* flagellar motor has 16 stator complexes, which form a large ring with 62 nm in diameter (Fig. 1**a, b**). Each stator complex includes a small, 8 nm ring within the cytoplasmic membrane (Fig. 1**a, b**). Improved resolution of the C-ring structure, obtained after focused refinement (Fig. 1**c** and Extended Data Fig. 3) shows 46-fold symmetry, consistent with that observed in the WT flagellar motors^18^.

**Figure 1.**
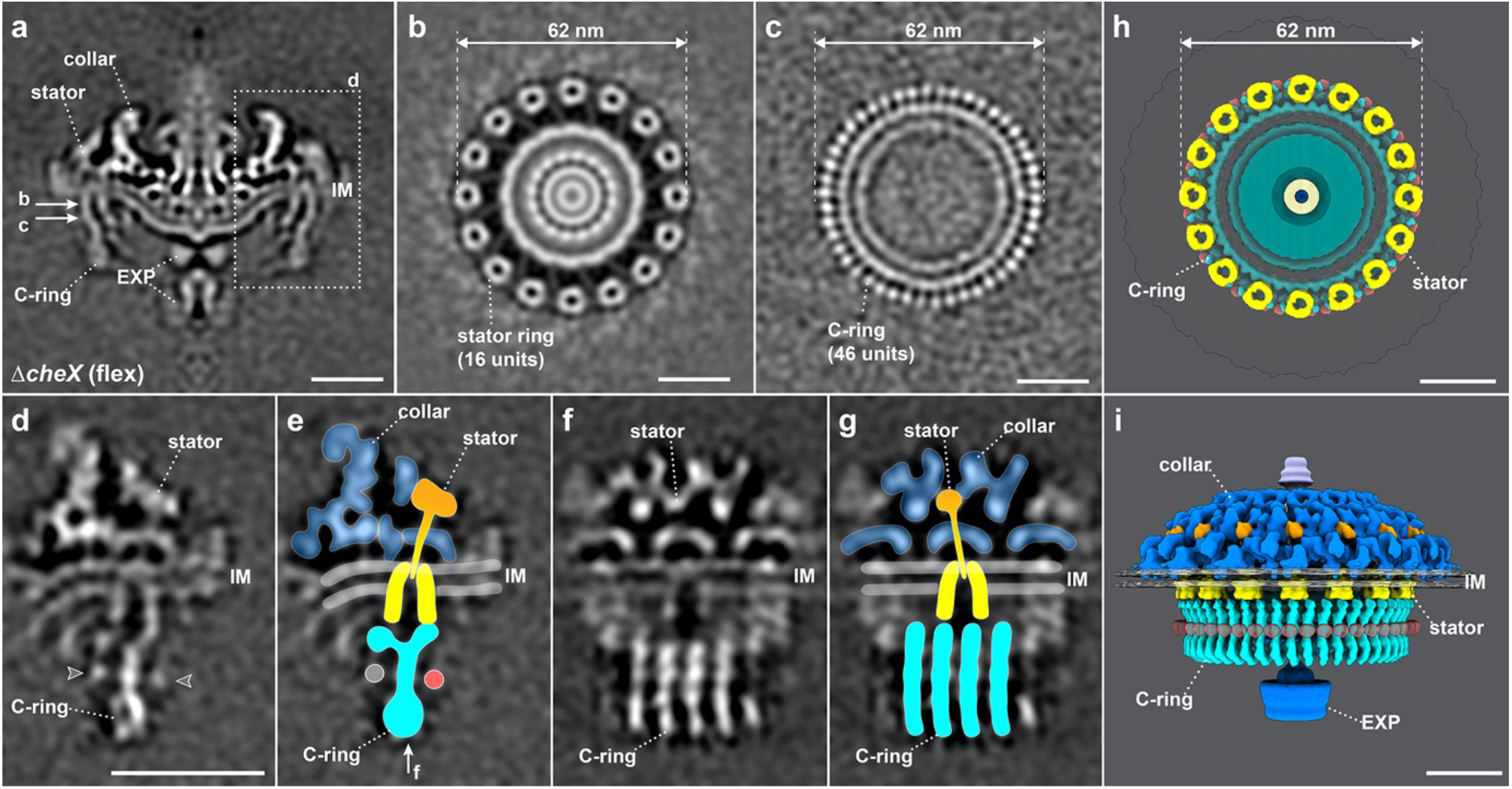
Structure of the flagellar motor in constantly flexing Δ*cheX* cells. (**a**) A medial cross-section of the *in situ* flagellar motor structure in Δ*cheX* determined by subtomogram averaging. The collar, stator, C-ring and export apparatus (EXP) are clearly visible in the cryoET map. (**b**) A perpendicular cross-section of the flagellar motor structure showing the stator ring. (**c**) The C-ring structure after focused alignment showing 46-fold symmetric features. (**d-g**) Stator-rotor interaction region (dash framed in panel **a**) after focused alignment. (**e, g**) The structures shown in (**d** and **f**) superimposed with the corresponding models in two different views. (**h**) A top view of the stator ring on the top of the C-ring. (**i**) A side view of the flagellar motor structure in 3D. Bar = 20 nm.

To further resolve detailed interaction between the C-ring and the stator complex, we applied symmetry expansion and utilized focused classification and alignment of the stator-rotor interaction region (dash framed region in Fig. 1**a**). The transmembrane and cytoplasmic portions of the stator complex have a bell-shaped structure embedded in the cytoplasmic membrane (Fig. 1**d-g**, Supplementary Video 2). It is 9 nm in height and 8 nm in diameter, which are similar to the dimensions of the purified MotA complex from *Aquafex aeolicus*^24^ (Extended Data Fig. 4). The periplasmic domain of the stator complex is inserted into the collar (Fig. 1**d-g**). It is ~9 nm long, and the top portion of its density corresponds well to the crystal structure of the *S. enterica* MotB periplasmic domain^25^ (Extended Data Fig. 4).

The C-ring exhibits a “Y” shape in the refined structure (Fig. 1**d, e**), which is similar to the previously reported *in vitro* C-ring structure in *S. enterica^26^.* However, the bottom portion of the C-ring in our structure is a spiral in which adjacent subunits are connected to one another. The top portion of the C-ring interacts with the periphery of the stator cytoplasmic region. By assembling the collar, stator complexes, and C-ring together, we revealed a complex architecture of the CW-rotating flagellar motor with unprecedented details (Fig. 1**h, i**).

### CheY3-P binds to the FliM protein of the C-ring

The well-defined C-ring in the Δ*cheX* mutant was found to be associated with two previously unidentified densities (arrowheads indicated in Fig. 1**d, e**). We hypothesized that these densities represent bound CheY3-P, as high levels of CheY3-P are expected in the Δ*cheX* cells^14^. To characterize CheY3-P and its interaction with the Δ*cheX* motor, we replaced the *cheX-cheY3* genes with *cheY3-gfp*, generating a *cheX::cheY3-GFP* mutant (Extended Data Fig. 5). Like Δ*cheX*, the GFP-labeled mutant constantly flexes. In addition, the mutant cells have fluorescent puncta at both cell poles (Fig. 2**a**), indicating that CheY3-P co-localizes with the flagellar motors. To confirm CheY3-P binding on the switch complex, we co-expressed His-CheY3 and FliM-FLAG in *E. coli* and affinity purified His-CheY3 and bound proteins by Ni-NTA binding in the presence or absence of acetyl phosphate (final concentration 40 mM). The purified products were examined using Western blots probed against anti-His or anti-FLAG antibodies. The His-CheY3* protein, in which Asp79 was converted to Ala, was used as a control, as it cannot be phosphorylated. In the presence of acetyl phosphate, FliM-FLAG co-purified with His-CheY3, but not with His-CheY3* (Fig. 2**e**, Extended Data Fig. 6). In the absence of acetyl phosphate, and therefore at low levels of CheY3-P, only a small amount of FliM-FLAG co-purified with His-CheY3. In contrast to FliM, no FliN-FLAG co-purified with His-CheY3, even in the presence of acetyl phosphate (Fig. 2**f**). These results indicate that CheY3 binds to FliM in a phosphorylation-dependent manner.

**Figure 2.**
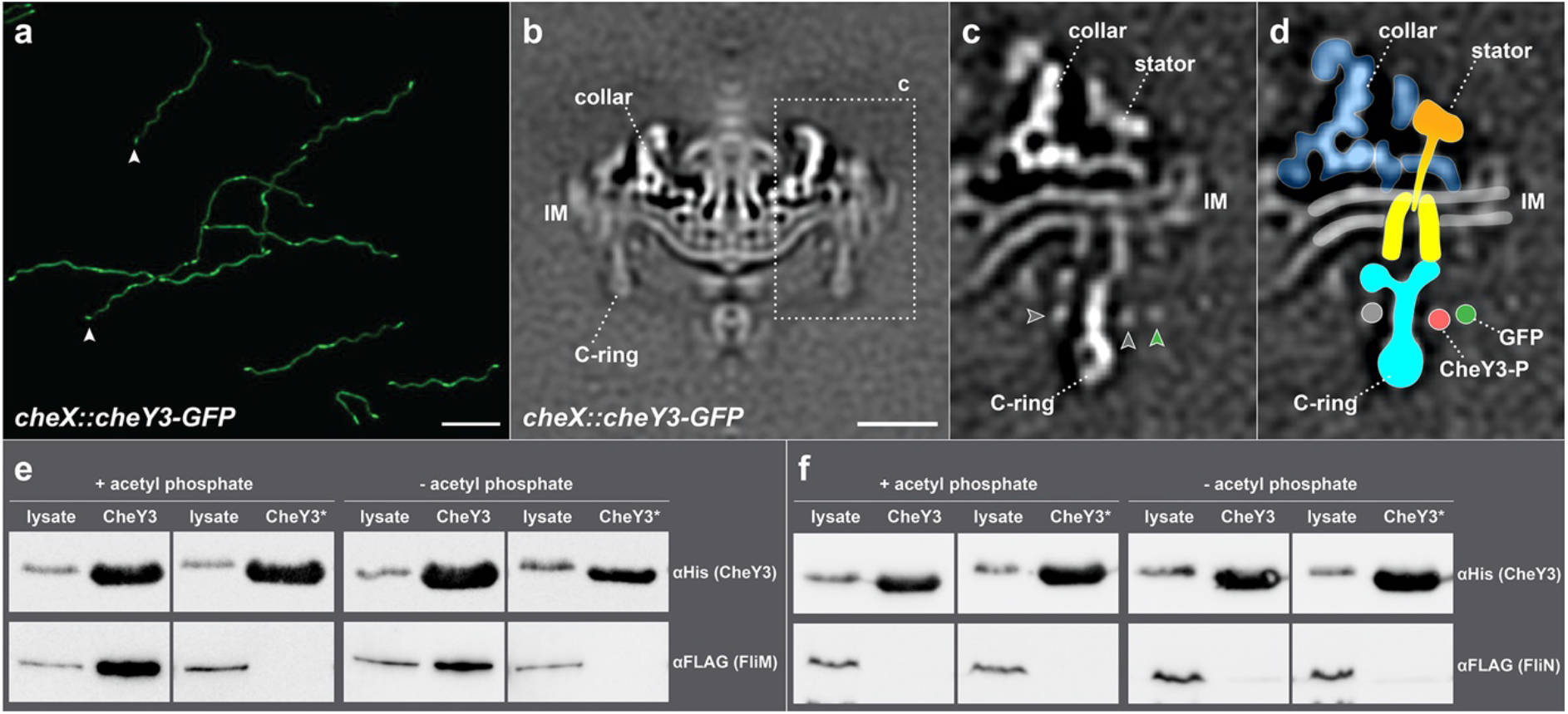
CheY3-P binding to the flagellar motor. (**a**) Fluorescence image of *cheX::cheY3-GFP* cells showing that GFP-tagged CheY3 proteins are polarly localized. (**b**) A medial cross-section of the flagellar motor structure in *cheX::cheY3-GFP* cells. (**c**) A refined structure of the stator-rotor interface (dash framed in panel **b**) in *cheX::cheY3-GFP.* Extra density (green arrow) is associated with the C-ring. (**d**) A cartoon model is superimposed onto the structure shown in panel **c**. The GFP density (green arrow indicated in panel **c** and green color highlighted in **d**) is located outside the C-ring. (**e, f**) Ni-NTA affinity purifications using the poly-histidine modified proteins HisCheY3 or HisCheY3* (CheY3^D79A^) to pull down FLAG-tagged FliM (FliM-FLAG) and FLAG-tagged FliN (FliN-FLAG), respectively. FliM-FLAG was co-purified with HisCheY3, but not with HisCheY3*, and more FliM-FLAG protein was co-purified with HisCheY3 in the presence of acetyl phosphate (**e**). There was no FliN-FLAG co-purified with HisCheY3/CheY3* (**f**). These results indicate that CheY3 binds to FliM protein in a phosphorylation-dependent manner. Bar = 10 μm in panel **a**, Bar = 20 nm in panel **b**.

To resolve the CheY3-P densities on the switch complex, we determined *in situ* structure of the motors in the *cheX::cheY3-GFP* mutant by cryoET and subtomogram averaging. Compared to the motor structure in the Δ*cheX* mutant, the motor structure in *cheX::cheY3-GFP* cells has an extra ring, likely contributed by GFP fused to CheY3 (green arrowhead in Fig. 2**c**). Together with the above biochemical data, we conclude that CheY3-P interacts with the FliM protein on the exterior side of the C-ring (Fig. 2**c, d**).

### Distinct conformations of the switch complex in the absence of CheY3-P

To compare the switch complex bound by CheY3-P with that in the absence of CheY3-P, we analyzed the motor structures in Δ*cheY3* mutant cells, which continuously run and cannot flex or reverse^27^ (Extended Data Table 1, Extended Data Fig. 2**b**, Supplementary Video 3). The overall *in situ* motor structure in the Δ*cheY3* mutant (Extended Data Fig. 7**a, b**) is quite similar to the averaged structure in the Δ*cheX* motor (Fig. 1**a, b**). Importantly, the stator ring is almost identical with 62nm in diameter. However, focused classification and alignment of the C-ring in the Δ*cheY3* motors revealed two distinct conformations of the C-ring: Δ*cheY3*-Class-1 (Fig. 3**a-e**, Extended Data Fig. 7**e-h**, Supplementary Video 4) and Δ*cheY3*-Class-2 (Fig. 3**f-j**, Extended Data Fig. 7**i-l**, Supplementary Video 5), although they share the same 46-fold symmetry and exhibit a “Y” shape structure. The C-ring conformation of Δ*cheY3*-Class-1 motors (Fig. 3**a-e**, Extended Data Fig. 7**h**) is similar to that in the Δ*cheX* motor (Fig. 1**d-g**, Extended Data Fig. 3**c**), while the CheY3-P density is absent. In contrast, the C-ring in Δ*cheY3*-Class-2 (Fig. 3**f-j**, Extended Data Fig. 7**i**) is twisted in a different direction compared to that in Δ*cheY3*-Class-1 (Fig. 3**a-e**, Extended Data Fig. 7**h**) or the Δ*cheX* motor (Fig. 1**d-g**, Extended Data Fig. 3**c**), resulting in different interactions between the stator and the C-ring. Specifically, the top portion of the C-ring in the Class-1 motor interacts with the outer part of the stator complex (Fig. 3**a, b**), which is the same as in the Δ*cheX* motor (Fig. 1**d, e**). In contrast, the top portion of the C-ring in the Class-2 motor interacts with the inner part of the stator complex (Fig. 3**f, g**). Therefore, our results suggest that the flagellar rotation direction is correlated with distinct stator-rotor interactions. As the Δ*cheY3* cells run constantly, we hypothesize that the Δ*cheY3*-Class-1 motors rotate CW near one pole, whereas the Δ*cheY3*-Class-2 motors rotate CCW near another pole. To test the model, we analyzed the motors at both poles in the same Δ*cheY3* cells. Our data confirmed that the motors near one pole indeed rotate CCW, while the motors near another pole in the same cell rotate CW (Extended Data Fig. 8).

**Figure 3.**
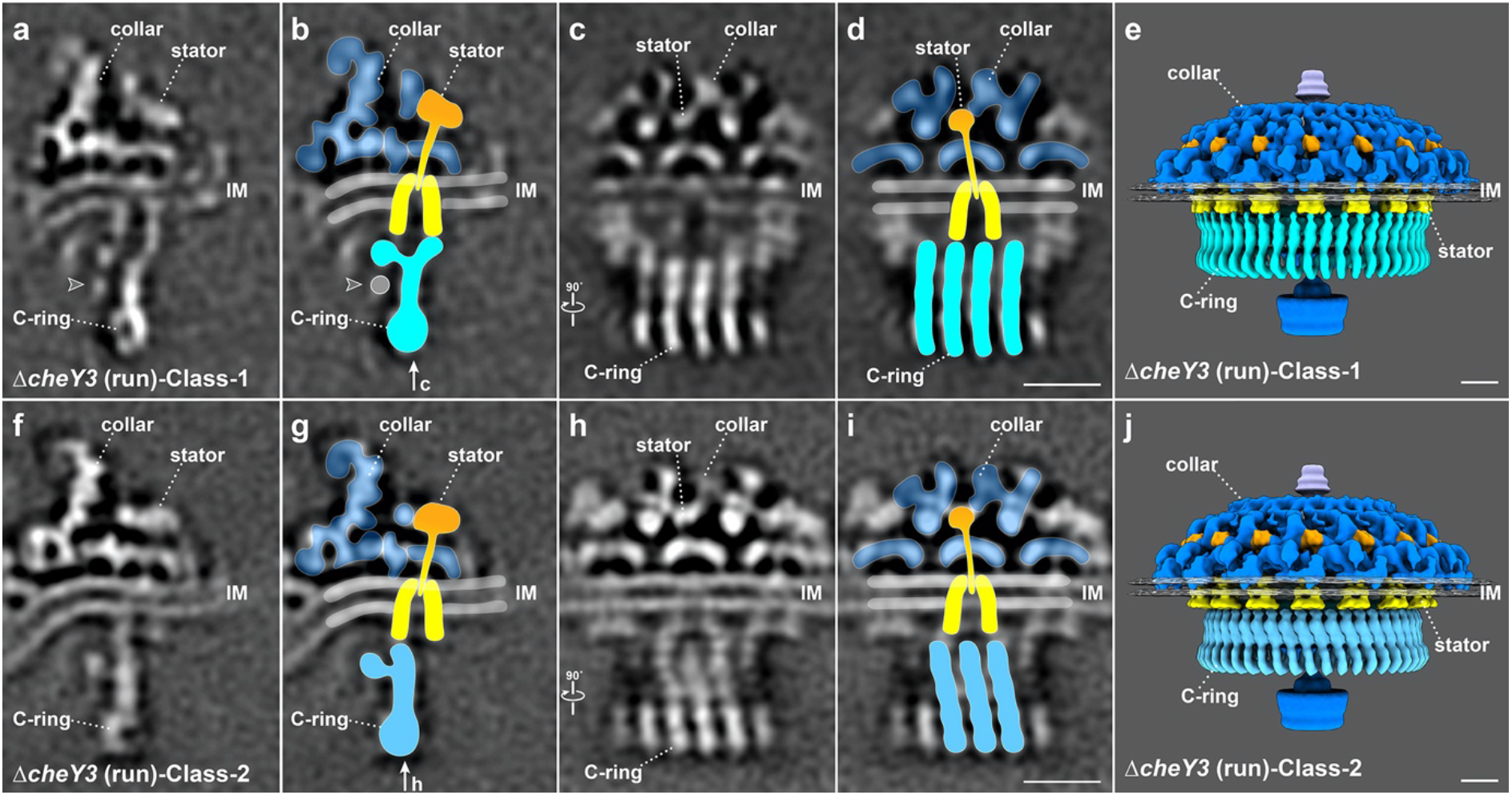
Stator-rotor interactions in constantly running Δ*cheY3* cells. Two distinct conformations of the C-ring are observed in Δ*cheY3* cells. (**a-e**) Detailed motor conformation in the Δ*cheY3*-Class-1 with the same views as shown in Fig.1**d-g, i**. (**f-j**) Detailed motor structures in the Δ*cheY3-Class-2.* The C-ring appears strikingly different in two class averages. In Class-1, the C-ring interacts with the outer part of the stator; while in Class-2, the C-ring interacts with the inner part of the stator. (**e, j**) 3D surface views of the Δ*cheY3-Class-1* and Δ*cheY3*-Class-2 flagellar motors. Note that the C-ring has two distinct conformations, enabling two different interactions with the stator complexes. Bar = 10 nm.

### CheY3-P binding triggers major remodeling of FliG2

To understand molecular details of the distinct C-ring conformations in the CW and CCW motors, we modeled the switch complex in the absence and presence of CheY3-P based on our cryoET maps and crystal structures of key flagellar components previously solved (see Methods). The resulting C-ring models fit well into our density maps (Fig. 4**b, f**). FliG2, a three-domain protein, forms the “v” at the top of the C-ring, poised to interact with the MS-ring via the N-terminal domain (FliG2_N_), and the stator complex via the C-terminal domain (FliG2_C_). The middle domain of FliG2 (FliG2_M_) interacts with the middle domain of FliM (FliM_M_), forming the stalk of the C-ring subunit. The C-terminal domain of FliM (FliM_C_) forms a heterodimer with FliN. A spiral is created at the base of the C-ring by alternating FliM_C_-FliN heterodimers and FliN-FliN homodimers (Fig. 4**c, d**), in which FliG2:FliM:FliN exist in a 1:1:3 stoichiometry as proposed previously^28,29^. The switch complex seen in the Δ*cheY3*-Class-2 motor represents the conformation associated with the CCW rotational state (Fig. 4**a-d**).

**Figure 4.**
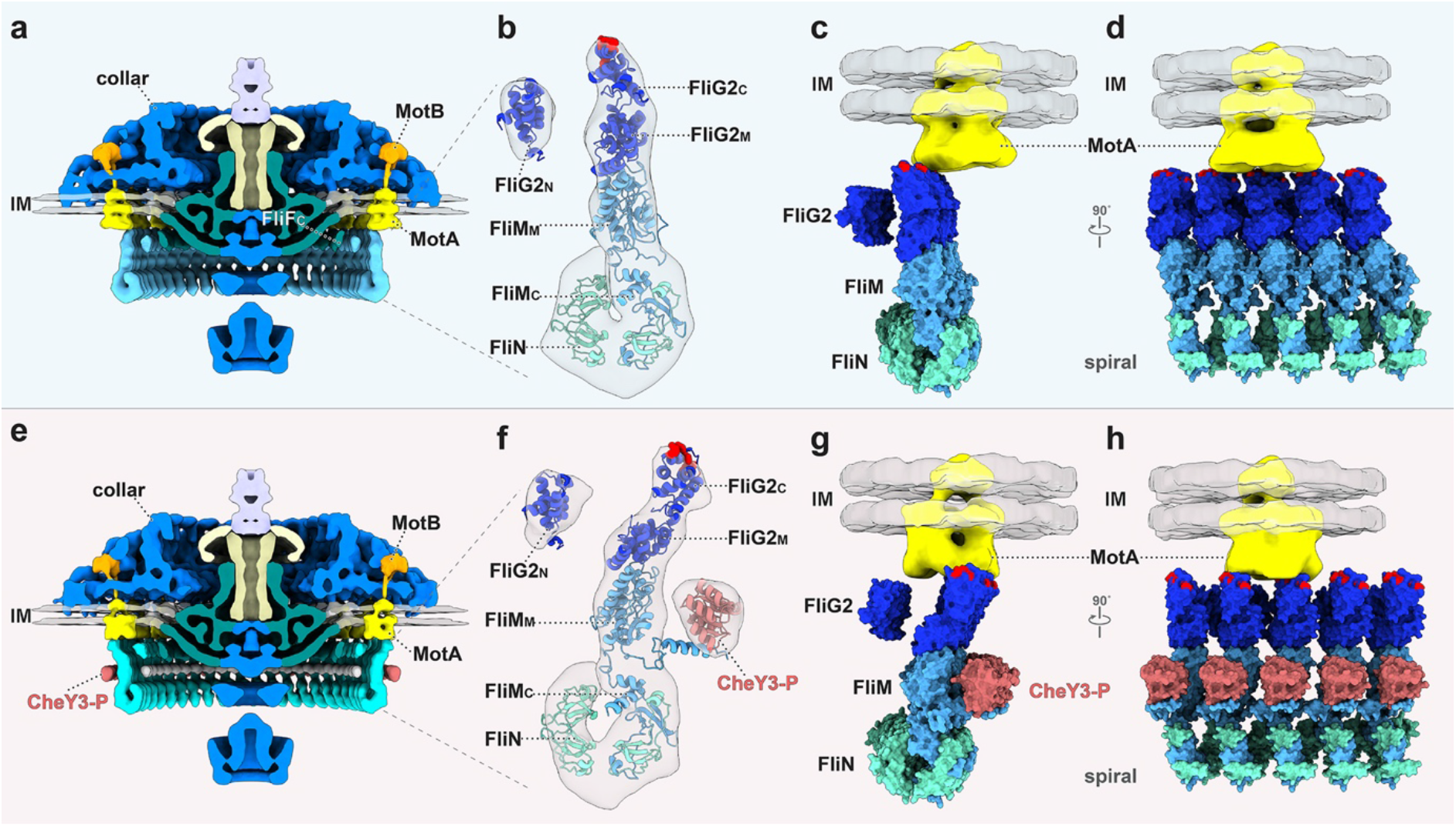
Molecular architectures of the flagellar motors without and with CheY3-P. (**a**) A medial cross-section of the flagellar motor structure without CheY3-P. (**b**) A pseudoatomic model of the C-ring unit shown in panel **a**. FliM and FliN have a stoichiometry of 1:3, and the C-terminal of FliM (FliM_C_) together with three FliN units form a spiral at the base of the C-ring. (**c**) Interactions between the bell-shaped stator complex and the C-ring. The charged residues (Lys275, Arg292, Glu299, and Asp300 in red) in FliG2_C_ interact with inner part of the stator complex. (**d**) A different view of five C-ring units connected at the based on the C-ring. (**e**) A medial cross-section of the flagellar motor structure in the presence of CheY3-P. (**f**) A pseudoatomic model of the C-ring unit with CheY3-P binding on the N-terminal of FliM (FliM_N_). (**g**) The charged residues (Lys275, Arg292, Glu299, and Asp300 in red) in FliG2_C_ interact with outer part of the stator complex. (**h**) A different view of four C-ring units are occupied by four CheY3-P proteins.

The switch complex of the Δ*cheX* motor is locked in the CW rotational state. When CheY3-P binds, the N-terminal domain of FliM (FliM_N_) interacts with CheY3-P, and this interaction results in an ~27° tilt of the FliM_M_ (Extended Data Fig. 9**b**). Importantly, although the spiral ring structure at the base of the C-ring remains almost the same, FliG2 undergoes a major remodeling in the Δ*cheX* motor (Fig. 4**e, f**) compared to that in Δ*cheY3*-Class-2 motor (Fig. 4**a, b**). The conformational change in FliG2 significantly enlarges the FliG2 ring from 55 nm to 62 nm, allowing FliG2 to interact with distinct parts of the stator ring (Fig. 4**c, g**, Extended Data Fig. 10).

## Discussion

Spirochetes have evolved a unique strategy to control motility^9,10^. However, it is still not clear how the WT spirochete produces asymmetric flagellar rotation. It is even more mysterious that asymmetric rotation persists in the complete absence of CheY3. In the constantly running Δ*cheY3* cells, we found two distinct conformations of the switch complex, consistent with the notion that they are in CW and CCW rotational states to keep the cell running (Extended Data Fig. 11). Comparison of the CW and CCW conformations in opposite poles of the constantly running Δ*cheY3* cells (Fig. 3**b, g**, Extended Data Fig. 8) reveals additional structure (colored in grey) associated with the C-ring in the CW conformation, suggesting that the additional structure likely plays a role in the asymmetric flagellar rotation in spirochete. As the extra structure and CheY3-P bind to FliM from two opposite sides of the C-ring, they may play similar roles in triggering the conformational change of FliG2 to allow CW rotation. The identity of the additional density is presently unknown. FliG1 is one possible candidate. It has been demonstrated previously that FliG2 is associated with the flagellar motors at both cell ends, whereas FliG1 is present at only one of the poles^23^. Δ*fliG2* cells are aflagellar and nonmotile, whereas Δ*fliG1* cells are flagellated, but have deficient motility in which the flagella at one cell pole appear to be ‘paralyzed’^23^. One possibility is that association of FliG1 with flagellar motors at one pole alters their ‘default’ structure and rotational direction, and their responses to regulatory elements such as CheY3-P. In addition, a double mutant lacking phosphodiesterases PdeA and PdeB has been shown to have a constantly flexing phenotype^30^, suggesting that these proteins might be also involved in regulating the asymmetric rotation. Further research is clearly required to clarify this issue.

Comparison of the CW motor in Δ*cheX* cells with the CCW Δ*cheY3*-Class-2 motor provides direct evidence for a profound conformational change in the C-ring caused by CheY3-P binding to FliM. The diameter of the FliG2 ring expands from 55 nm to 62 nm upon binding of CheY3-P, whereas the diameter of the bottom portion of the C-ring remains similar (Extended Data Fig. 10). Importantly, the *B. burgdorferi* C-ring in both CCW and CW rotations possesses 46-fold symmetry, with each unit composed of FliG2, FliM, and FliN (1:1:3). In each subunit, one FliM and three FliN proteins form the base in a spiral shape, and one FliG2 stacks on FliM. The dramatic conformational changes in FliG2 are accommodated by the flexibility of the helical linker between the FliG2_MC_ domains^31-33^. This helix contains a highly conserved Gly-Gly residue pair located near the C-terminus of the helix^32,34,35^. The large rearrangement of FliG2 during directional switching allows it to engage different parts of the stator complex in the CW and CCW conformations.

The stator complex is known to be highly dynamic in many bacterial species^25,36^. As a result, it has been very challenging to visualize the stator complex in the intact motor at high resolution^15,16,18^. Here, we used cryoET and focused refinement to visualize the bell-shaped structure of the stator complex in both the CW and CCW rotational states. This finding is of particular importance, because it allows us to understand how the stator complex interacts with the switch complex at the molecular level. Sixteen bell-shaped stator complexes form a stator ring of 62 nm in diameter. In CW rotation, FliG2_C_ interacts with the outer part of the stator ring, while during CCW rotation it interacts with the inner part of the stator ring. This association suggests that the outer part of stator cytoplasmic region drives the C-ring CW, while the inner part drives the C-ring CCW. This result is consistent with the notion that the inward flow of protons drives the unidirectional rotation of the MotA portion of the stator. Based on these predictions, we propose a novel model for the generation of flagellar rotation and for the switching of rotational directions (Fig. 5, Supplementary Video 6).

**Figure 5.**
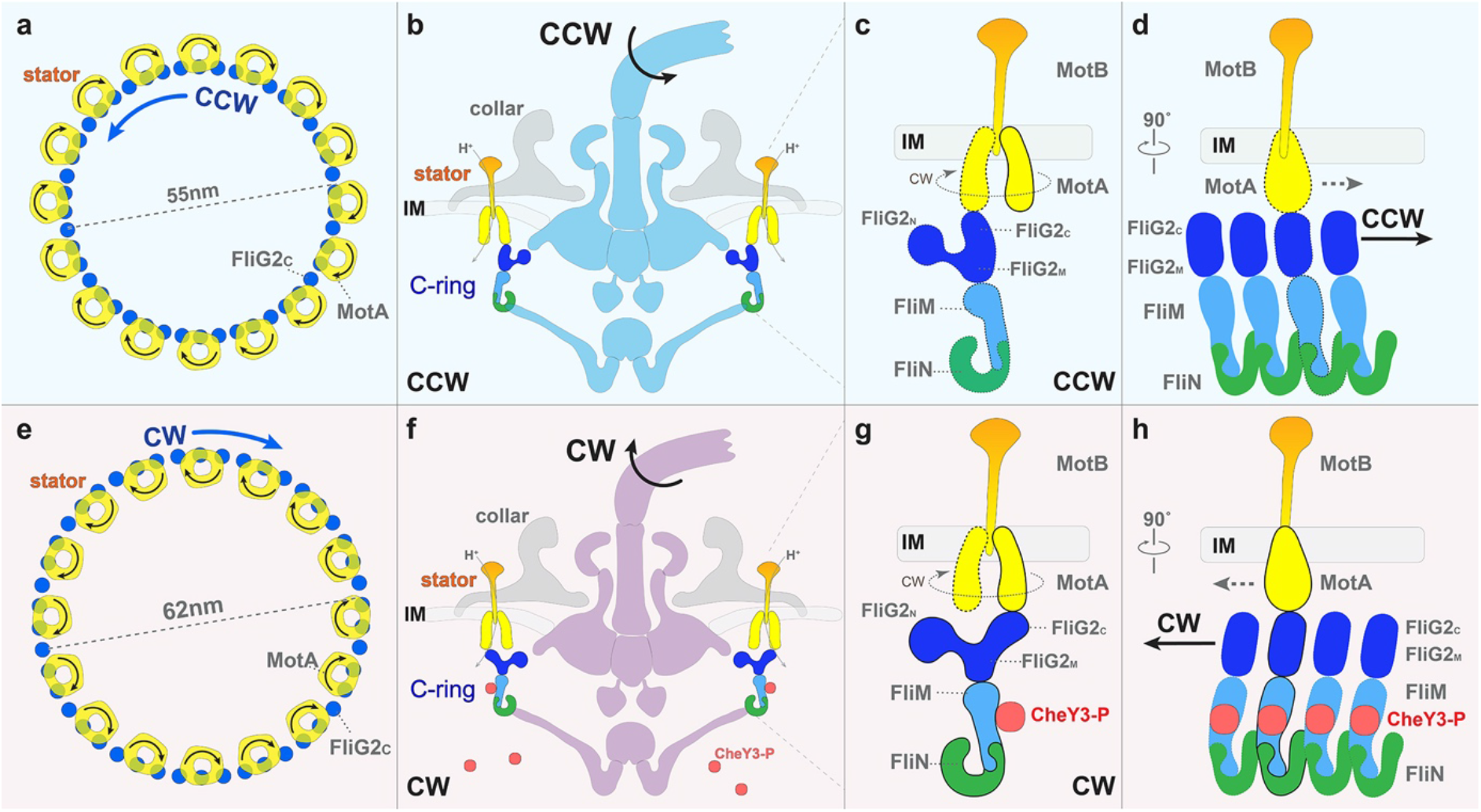
Model for the mechanism of rotational switching. (**a, b**) Interactions of the stator with FliG2 in the C-ring during CCW rotation. In the default state when there is no bound CheY3-P, the FliG2 proteins interact with the inner part of the stator complex (colored in yellow). With the influx of protons through the stator channel, the cytoplasmic subunits of each stator complex spins CW. Therefore, the C-ring (blue) is induced to spin CCW. (**c**) A zoomed-in view of the interaction between the C-ring and the stator complex. (**d**) A perpendicular view shows that four C-ring units are connected by FliM/FliN interactions. (**e, f**) CheY3-P induced conformational changes in the C-ring result in altered interactions between the stator and C-ring, thereby causing the switch to CW rotation. When CheY3-P binds to FliM on the exterior surface of the C-ring, its binding triggers the shift (**g**) and tilt (**h**) of FliG2 so that FliG2_C_ interacts with the outer part of the cytoplasmic domain of the stator complex (**g**). Because the cytoplasmic domain of the stator always spins CW, the C-ring is induced to spin CW (**e**). During the rotational switch, the spiral ring structure formed by FliM and FliN acts as a base to hold the C-ring structure together (**d, h**).

When protons flow inward through the stator ion channel, we postulate that the cytoplasmic region of the stator rotates CW (viewing from MotB through the membrane to MotA). In the default state (without CheY3-P) FliG2 interacts with the inner part of the stator cytoplasmic region and the C-ring rotates CCW (Fig. 5**a, b**). When CheY3-P binds to FliM from the exterior side of the C-ring (Fig. 5**f, g**), FliG2 undergoes a major remodeling to interact the outer part of the stator (Fig. 5**f, g**). The interaction with the CW-rotating stator would then drive CW rotation of the C-ring (Fig. 5**e**). As FliM and FliN form a stable spiral ring at the base of the C-ring (Fig. 5**d, h**), the CheY3-P mediated conformational changes of FliG2 allow rapid rotational switching. Given that the C-ring and stator are evolutionarily conserved, this molecular mechanism for flagellar rotational switching may be utilized, with some modifications, across a wide spectrum of bacterial species.

Many challenges remain to test this model. The most obvious one is to directly demonstrate that the cytoplasmic domains of the stator units actually rotate, although a recent study on Tom complex, a homologous complex of the stator complex, suggested that it may form a pentamer and rotate in presence of the proton motive force^37^. Each stator unit contains a central MotB dimer and four to five peripheral MotA subunits. MotB is stationary; in *B. burgdorferi* it is embedded in the collar and firmly attached to the peptidoglycan of the cell wall. The critical conserved Asp residue required for proton conduction is on the single transmembrane helix of MotB^17^. The model predicts that the MotA subunits rotate around MotB in a manner that is coupled to the inward flow of protons, resulting in sequential interactions of the MotA subunits with consecutive FliG2 units in the C-ring (Fig. 5). It must be remembered that the transmembrane helices of MotA and MotB are close together at the base of the splayed bell-shaped structure of the stator cytoplasmic domain.

In summary, we determined the structures of CW- and CCW-rotating flagellar motors in *B. burgdorferi* by cryoET and subtomogram averaging. We demonstrated that the flagellar switch complexes undergo substantial remodeling to form distinct interactions with the stator complexes during the rotational switching, analogous to throwing an automobile transmission into reverse. We propose a novel model for the generation of torque and the switching of rotational direction. A proton flux through the stator causes the bell-shaped MotA cytoplasmic region to rotate CW (view from the hook to the C-ring). Interactions with the outer part of the stator cytoplasmic region cause the C-ring to rotate CW, and interactions with the inner part of the stator cytoplasmic region cause the C-ring to rotate CCW. Control of the direction of flagellar rotation consists of aligning the interaction sites of the stator and the switch complex properly through conformational changes in FliG2 to achieve the desired direction of flagellar rotation.

## Acknowledgements

We thank Michael Manson, Justin Radolf, and Michio Homma for critical reading and suggestion. This work was supported by grants from the National Institute of Allergy and Infectious Diseases (R01AI087946, R01AI078958, R01AI132818, and R01AI59048) and the National Institute of Dental and Craniofacial Research (R01DE023080). This work is dedicated to the memory of Prof. Fanghua Li, who was an incredible scientist and a trusted mentor for YC, JL, and many others in the field of electron microscopy.

## Author contributions

J.L. and C.L. conceived the project. Y.C. performed cryoET experiments, data analysis, modeling and wrote the manuscript draft. K.Z. performed genetical and biochemical experiments and analysis. B.C. and X. Z. contributed structural analysis. J.L. and C. L. supervised all work. M.M., S.J.N, and N.W.C. provided *B. burgdorferi* strains, Y.C, C.L., and J.L. prepared the manuscript with input from all authors.

## Materials and Methods

### Bacterial strains and growth conditions

A high-passage *B. burgdorferi* sensu stricto strain B31A (WT) and its isogenic mutants were grown in Barbour-Stoenner-Kelly II (BSK-II) liquid medium or on semisolid agar plates at 34°C in the presence of 3.4% carbon dioxide as previously described^38,39^.

*Escherichia coli* TOP10 strain (Invitrogen, Carlsbad, CA, USA) was used for DNA cloning and plasmid amplifications. BL21 strains transformed with GroEL-GroES chaperones (Takara Bio USA) were used for recombinant protein preparations. *E. coli* strains were cultured in lysogeny broth (LB) supplemented with appropriate antibiotics as needed. The Δ*cheX* and Δ*cheY3* mutants of *B. burgdorferi* were constructed and characterized as previously described^40,41^.

### Inactivation of *cheX* using *cheY3-gfp*

The vectors for in frame replacing *cheX-cheY3* with *cheY3-gfp* were constructed by using a PCR-based fusion method as previously described^41^. Briefly, the PCR primers (containing complementary overlaps to downstream fragment) were designed immediately flanking the *cheX-cheY3* genes, to generate approximate 1-kb products upstream and downstream of the coding sequences. The primers for the *flgB* promoter, *cheY3, gfp* and streptomycin resistance cassette *(str)* were designed. Initial PCR amplifications for each of individual fragments (i.e., 5’- and 3’-flanking DNA of *cheX-cheY3, flgB* promoter, *cheY3, gfp*, and *str*) were performed, followed by a fusion PCR connecting all the fragments together, generating the constructs of *cheY3-gfp::str* (Extended Data Fig. 5). The resultant constructs were transformed into competent B31A cells by electroporation to delete *cheX-cheY3* genes. The resultant mutant clones were confirmed by PCR and western blots using antibodies against GFP, CheY3 and CheX, respectively.

### Light and fluorescence Microscopy

Fluorescence images of *cheX::cheY3-GFP B. burgdorferi* cells were taken using a Zeiss Axiostar plus microscope at a wavelength of 480 nm. The images were captured and processed using the program ZEN (Zeiss, Germany).

### Co-expression and purification of CheY3/CheY3* and FliM/FliN

The full-length *cheY3* or *cheY3*\* (in *cheY3*\*, Asp79, a key residue required for the phosphorylation, was replaced by Ala (Extended Data Table 2). Che *cheY3* gene was first amplified by PCR (primers P15/P16) using DNA polymerase (Invitrogen, Carlsbad, CA) with engineered BamHI and SacI cut sites at its 5′ and 3′ ends, respectively. The amplicon was cloned into the pGEM-T Easy vector (Promega, Madison, WI) and then subcloned into the pQE80 expression vector (Qiagen, Valencia, CA), yielding a vector of pQE80CheY3/CheY3*, and thereby incorporating an N-terminal histidine (His) tag. The full-length *fliM and fliN* genes (without stop codon) were PCR amplified (primers P17/P18 and P19/P20) using DNA polymerase (Invitrogen, Carlsbad, CA) with engineered SacI at its 5′ end and FLAG tag and SalI cut site at the 3′ ends. The amplicons were first cloned into the pGEM-T Easy vector and then digested using SacI and SalI and subcloned into precut pQE80CheY3/CheY3*. The resultant plasmid was then transformed into the BL21 strain that harbors GroEL-GroES chaperones for protein production. The expression of recombinant proteins in *E. coli* cells was induced with 1 mM isopropyl-β-D-thiogalactoside (IPTG) overnight at 16°C. Recombinant HisCheY or HisCheY* and bound proteins were purified using nickel agarose columns (Qiagen) under native conditions per manufacturers’ instructions.

### Site-directed mutagenesis of CheY3

Site-directed mutagenesis was performed using QuikChange site-directed mutagenesis kit (Stratagene, San Diego, CA) per manufacturer’s instructions. The above constructed cheY3 pGEM-T Easy vector was used as a template for the mutagenesis. Amino acids in CheY3 (Asp79) were substituted with Ala, using primers P13/P14. The mutation was confirmed by DNA sequencing analysis. The mutated genes were PCR amplified and subcloned into pQE80 expression vector as described before.

### Preparation of cryoET samples

*B. burgdorferi* cells were cultured to log phase, then centrifuged in 1.5 ml tubes at 4-5,000 rpm for ~5 minutes, the resulting pellet was rinsed gently with 1 mL phosphate buffered saline (PBS). The cells were centrifuged again and finally suspended in 20-100 μl PBS to obtain an appropriate concentration for plunge freezing. The cell solution was then mixed with 10 nm colloidal gold fiducial markers. CryoET samples were prepared using copper grids with holey carbon support film (200 mesh, R2/1, Quantifoil). The grids were glow-discharged for 30 seconds before we deposited 5 μL cell solution on them. Then the grids were blotted with filter paper and rapidly frozen in liquid ethane using a homemade gravity-driven plunger apparatus.

### CryoET data collection and tomogram reconstruction

The frozen-hydrated samples were transferred to a 300 kV Titan Krios electron microscope (Thermo Fisher) equipped with a Direct Electron Detector and energy filter (Gatan). For the Δ*cheX* and Δ*cheY3* samples: Images were recorded at 53K magnification with pixel size of 2.747 Å. SerialEM^42^ was used to collect tilt series at 2 to 4 μm defocus, starting from 36°, with accumulative dose of ~60-70 e^-^/Å^2^ distributed over 35 images and covering angles from −51°to 51°, with a tilt step of 3°. For the *cheX::cheY3-GFP* sample: Images were recorded at 64K magnification with pixel size of 2.245 Å. The tilt series were collected in two different strategies using SerialEM^42^ with accumulated dose ~80 e^-^/Å^2^. Strategy 1: tilt series were collected under super-resolution, with 2 to 4 μm defocus using the implemented dose-symmetric tilt scheme in SerialEM^42^. The dose-symmetric tilt scheme parameters were set as: start from 0°, tilt from −51° to 51° with 3° tilt step, group size 2 and stop alternating directions beyond 36° from the initial angle. Strategy 2: tilt series were collected using the improved Fast Incremental Single Exposure method^43^ with the dose-symmetric tilt scheme, 2 to 4 μm defocus, tilt from −54° to 54° with 3° tilt step and group size 2. 133 and 56 tilt series were collected in strategy 1 and strategy 2, respectively.

All recorded images were first motion-corrected using MotionCorr2^44^ and then stacked by IMOD^45^. The tilt series were aligned using fiducial markers or fiducial free alignment by IMOD. Gctf^46^ was used to determine the defocus of each tilt image in the aligned stacks and the “ctfphaseflip” function in IMOD was used to do the contrast transfer function (CTF) correction for the tilt images. Tomograms were then reconstructed by weighted back-projection using IMOD^45^ with the CTF corrected aligned stacks.

### Subtomogram averaging and corresponding analysis

Bacterial flagellar motors were manually picked from the bin6 tomograms as described^47^. The subtomograms of flagellar motors were first extracted from the bin6 tomograms, then the i3 software package^48,49^ was used for 3D alignment and classification to get the refined particle positions and remove junk particles. Afterwards, the subtomograms were extracted from unbinned tomograms with the refined positions and furtherly binned by 2 or 4 based on the requirement for alignment and classification. In total, 1,065 subtomograms of Δ*cheX* motors, 2,087 subtomograms of Δ*cheY3* motors and 1,250 subtomograms of *cheX::cheY3-GFP* motors (879 motors and 371 motors from tilt series collected in strategy 1 and strategy 2, respectively) were selected from the tomographic reconstructions and used for subtomogram analysis. Class averages were computed in Fourier space so that the missing wedge problem of tomography was minimized^49,50^. Gold standard Fourier shell correlation coefficients were calculated by generating the correlation between two randomly divided halves of the aligned images used to estimate the resolution and to generate the final maps.

Focused refinement, multi-reference alignment (MRA) and 3D classification of the whole C-ring structure: after we got the initial whole motor structure, we did small angular search along the motor rod to refine the C-ring structure. During the refinement, a 3D molecular mask slightly bigger than the C-ring part was applied to the reference and the angular search range was restricted to be smaller than ±5° so that we can maintain the overall alignment of the motor. Then we got the refined C-ring structures in Δ*cheX* (Extended Data Fig. 3c) and Δ*cheY3* (Extended Data Fig. 7d) flagellar motors after several cycles’ refinement. Afterwards, these two C-ring structures were used as the references for the MRA. MRA was applied for both Δ*cheX* and Δ*cheY3* mutants followed by 3D classification. Then we got the two different C-ring conformations in Δ*cheY3* motors (Extended Data Fig. 7h, l), but just one C-ring conformation in Δ*cheX* motors.

Focused refinement of the stator-rotor interaction region: each flagellar motor has 16 stator complexes. After the alignment for the whole motor structure, the regions around 16 stator complexes were first extracted from each motor, then we refined the 3D alignment and applied 3D classification to remove particles with bad contrast or large distortions to get the refined structures. Such focused refinement was applied to four motor sets: the Δ*cheX* motors, the Δ*cheY3*-Class1 motors, the Δ*cheY3*-Class2 motors and the *cheX::cheY3-GFP* motors.

Focused alignment of the C-ring subunit in different cell tips of Δ*cheY3* cells (Extended Data Fig. 8): for the motors at one tip of a Δ*cheY3* cell, we first aligned the whole motor structure, then we applied symmetry expansion based on the C-ring symmetry (46-fold symmetry) and did MRA alignment for the C-ring subunit (dash framed region in (Extended Data Fig. 8**d**) to generate a refined structure (Extended Data Fig. 8**e, f**)). The C-ring structures shown in (Extended Data Fig. 7**h, l**) were used as references for the MRA alignment. Afterwards, we can identify the rotation direction of these motors based on the twist direction of the refined C-ring subunit. The rotation direction of the motors at another cell tip were determined in the same way. Similar analysis was applied to other 4 Δ*cheY3* cells. The motors at different cell tips were found to rotate in opposite directions, then we merged all CCW or CW rotating motors together and generated the structures shown in (Extended Data Fig. 8**j-l**) or (Extended Data Fig. 8**m-o**), respectively.

### Model generation and refinement

Based on the reported crystal structures of FliG_C_ and FliG_M_ (PDB 4FHR)^51^, FliG_N_ (PDB 3HJL)^52^, FliM_C_ and FliM_M_ (PDB 4FHR)^51^, FliM_N_ (PDB 4YXB)^53^, FliN (PDB 1YAB)^54^, and CheY (PDB 4IGA)^55^, the *B. burgdorferi* C-ring proteins were generated using I-TASSER^56-58^. FliG2, FliM, and FliN were placed into the cryoET maps by using UCSF Chimera^59^. The unknown protein-protein interfaces were refined in Rosetta using the protein-protein docking scripts^60^. The model was refined using PHENIX Real Space Refinement^61^ to move the protein domains relative to one another while preserving the known architecture of the C-ring subunits. MotB (PDB 2ZVY)^62^ and MotA (EMD 3417)^63^ were used to fit into the cryoET maps by using UCSF Chimera^59^.

### Three-dimensional visualization

UCSF Chimera^59^ and UCSF ChimeraX^64^ were used for surface rendering of subtomogram averages, segmentation, and molecular modeling. Unrolled maps of the motor structures were generated using ‘vop unroll’ function of UCSF Chimera^59^.

## Extended Data Figures and Tables

**Extended Data Fig. 1.**
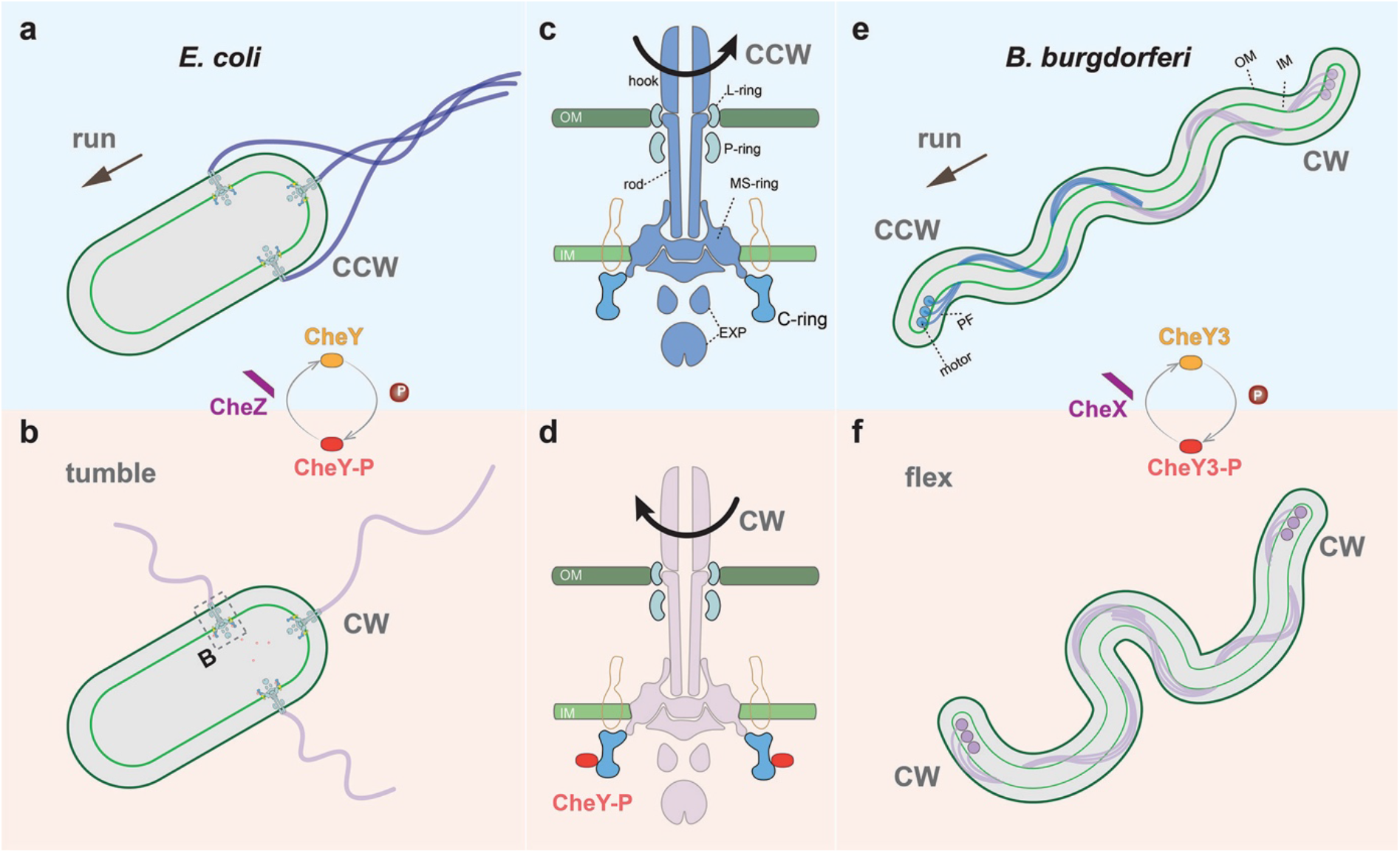
Swimming motility modes and flagellar switching in *E. coli* and *B. burgdorferi*. (**a, b**) Cartoon of the swimming motility modes in *E. coli:* run and tumble. (**c**) The motor rotates CCW as a default state. (**d**) When the level of CheY-P becomes high enough, CheY-P binds to the C-ring, and the motor switches to CW rotation. The chemotaxis protein CheZ dephosphorylates CheY-P to return the motor to CCW rotation. (**e, f**) Swimming motility modes in *B. burgdorferi*: run and flex. Periplasmic flagella (PF) are located between the inner membrane (IM) and outer membrane (OM). The flagellar motors are attached near each cell pole. Spirochetes run when the anterior flagella rotate CCW and the posterior flagella rotate CW (**e**). When the flagella at both poles rotate in the same direction (CW), the spirochetes flex in place and fail to move translationally. The swimming motility of *B. burgdorferi* is also controlled by a chemotaxis system. The homologs of CheY and CheZ in *B. burgdorferi* are CheY3 and CheX.

**Extended Data Fig. 2.**
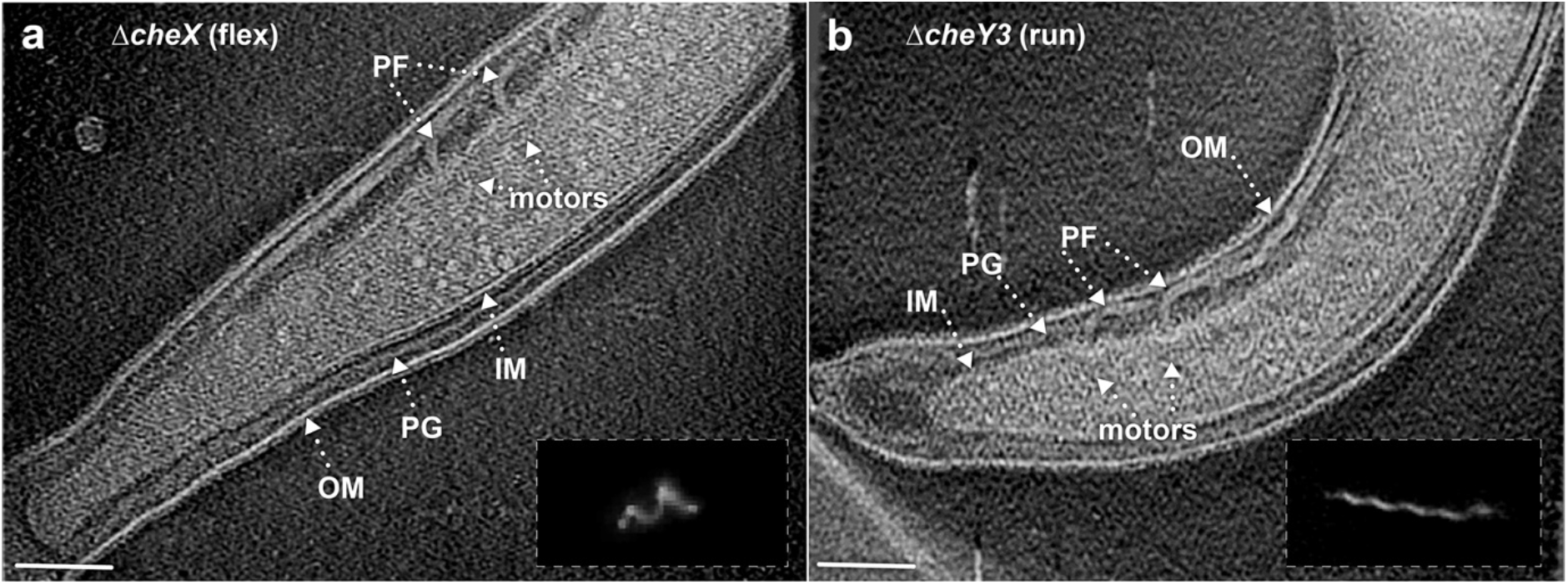
Cryo-ET imaging of the flagellar motors in Δ*cheX* and Δ*cheY3* mutants. (**a**) A representative tomographic section from a Δ*cheX* cell tip reconstruction. Outer membrane (OM), inner membrane (IM), peptidoglycan layer (PG), and motors are clearly resolved in the tomogram. (**b**) A representative section of a tomogram from a Δ*cheY3* cell tip. Multiple motors with different orientations can be found at the cell tip. The insertions in (**a, b**) are the dark-field images showing a Δ*cheX* cell constantly flexing and a constantly running Δ*cheY3* cell, respectively.

**Extended Data Fig. 3.**
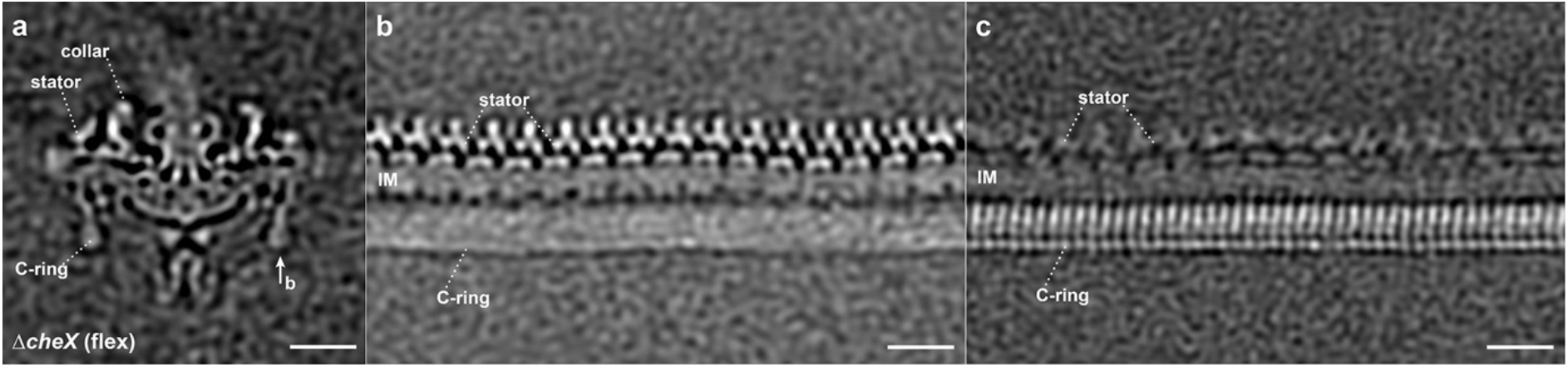
Refined structure of the C-ring in the Δ*cheX* motor. (**a**) A medial cross-section of an averaged map of the Δ*cheX* motor. (**b**) The unrolled map refined using the stator region densities shows 16 stator complexes are embedded in the inner membrane (IM), while the C-ring subunits are unresolved due to symmetry mismatch between the C-ring and the stator. (**c**) The unrolled map refined using the C-ring region densities shows 46-fold symmetric features, while the stator becomes blurry. Bar = 20 nm.

**Extended Data Fig. 4.**
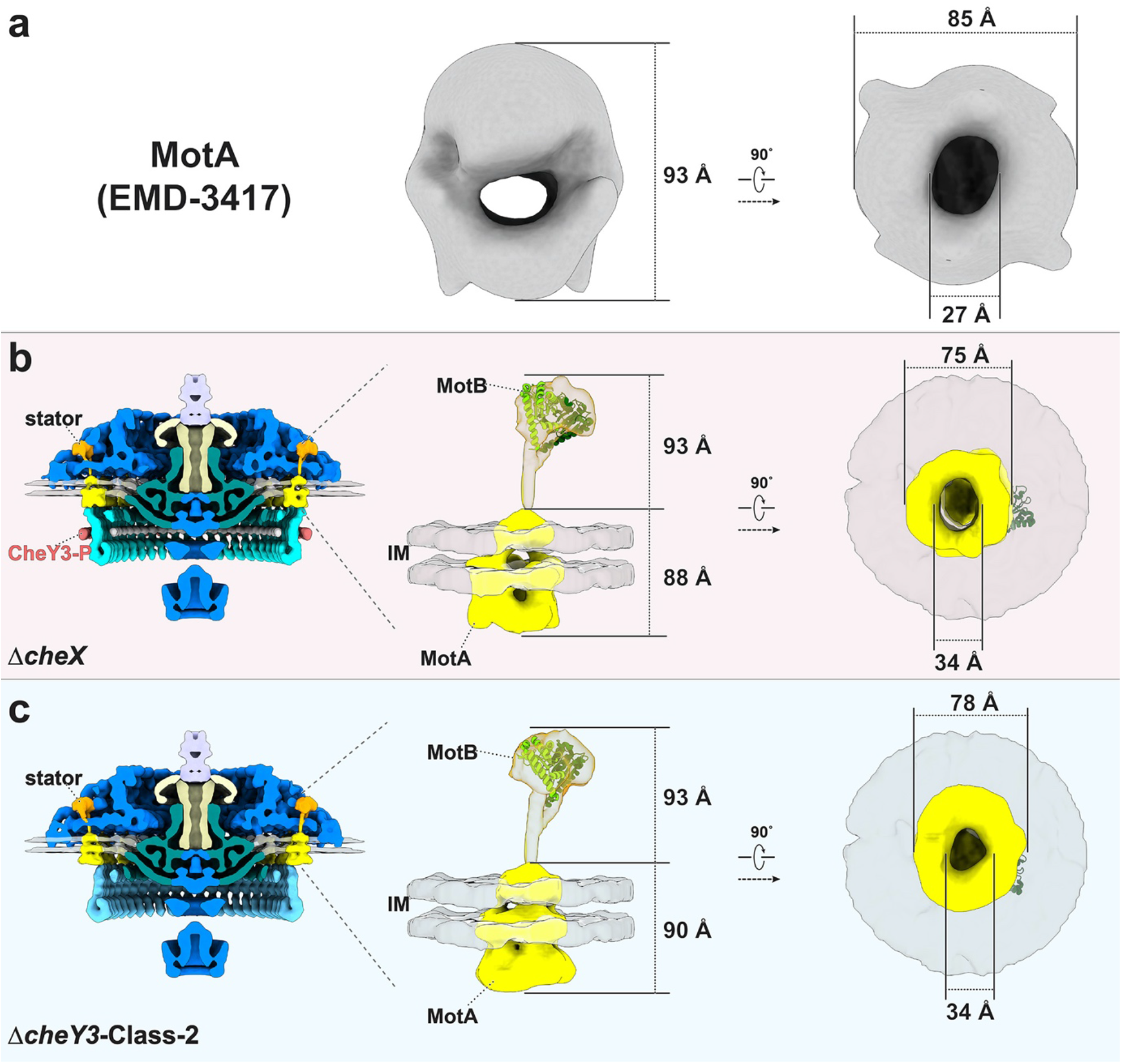
Comparison between *in situ* stator complex and the purified stator components. (**a**) Structure of purified MotA complex from *A. aeolicus* resolved by single particle EM (EMD 3417)^63^. (**b**) The *in situ* stator complex in the Δ*cheX* motor has a bell-shaped structure embedded in the inner membrane (IM) and a periplasmic domain. The top part of the periplasmic domain matches well with the crystal structure of the *S. enterica* MotB periplasmic domain (PDB 2ZVY)^62^ (middle panel). The bell-shaped structure has similar size and shape as the structure of EMD 3417 (right panel). (**c**) The *in situ* stator complex in the Δ*cheY3-Class-2* motor is similar to that in the Δ*cheX* motor.

**Extended Data Fig. 5.**
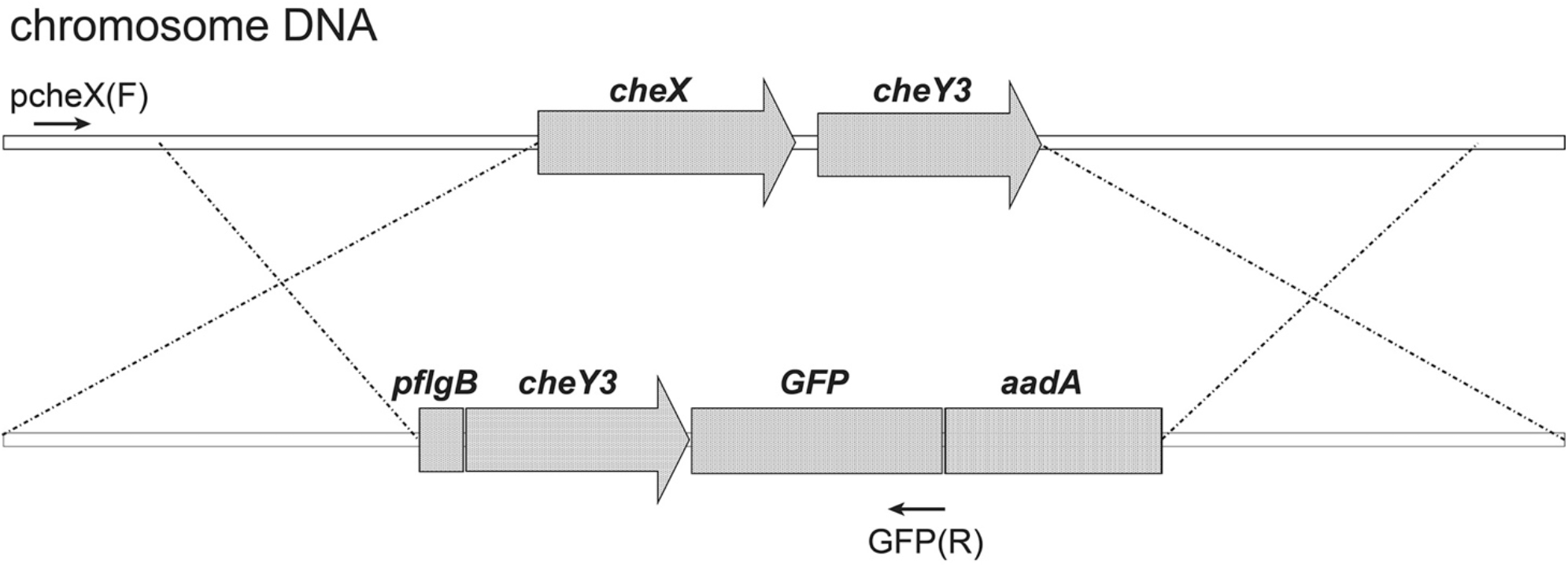
Schematic diagram for the in-frame replacement of *cheX-cheY3* genes with *cheY3-gfp*. *aadA*, a streptomycin resistance gene was used as a selection marker. pcheX(F) and GFP (R) are oligonucleotide primers utilized to verify the occurrence of the allelic exchange of the recombinant construct (bottom) into the targeted region in the *B. burgdorferi* chromosome (top).

**Extended Data Fig. 6.**
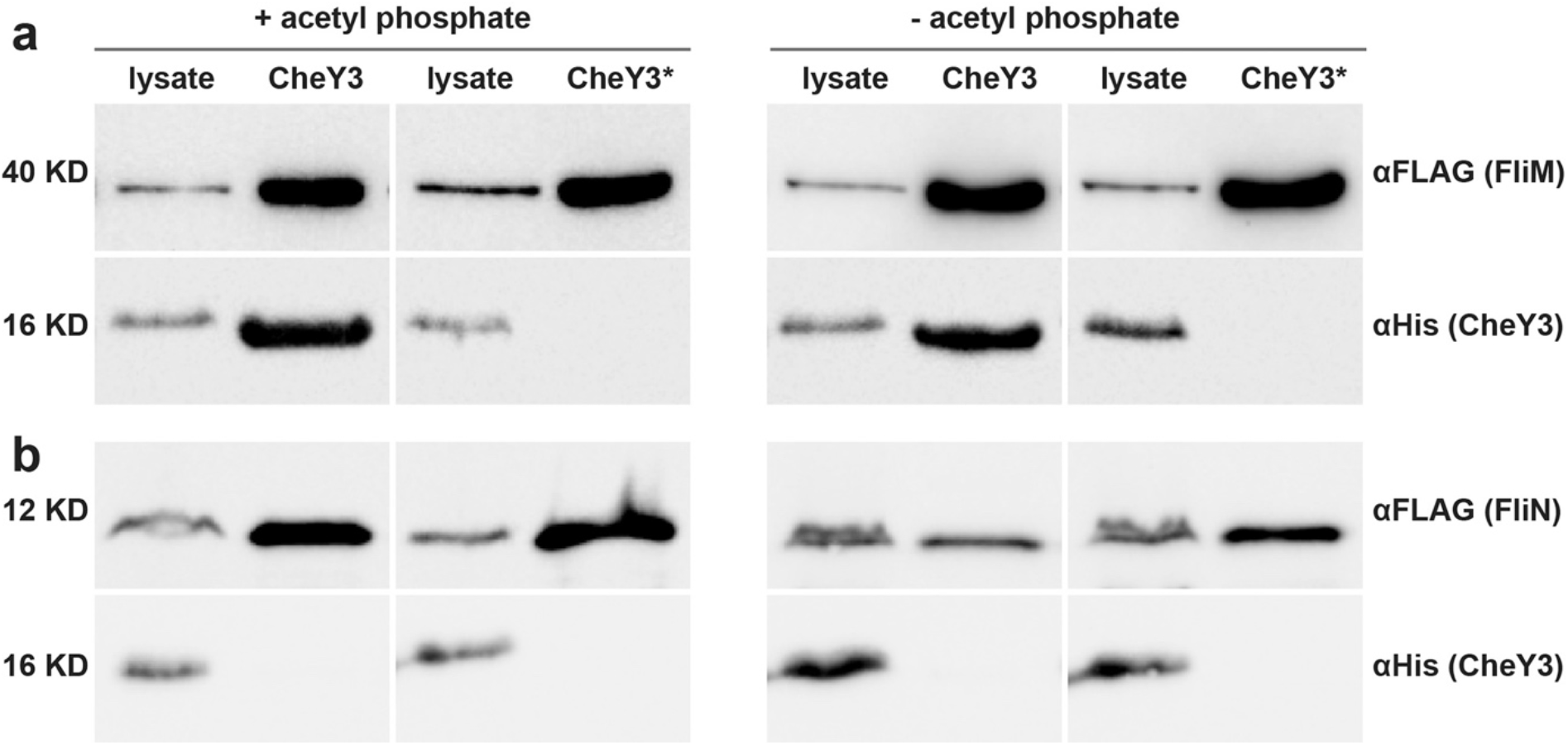
FLAG affinity purification using FLAG-FliM/FliN to pull down HisCheY3/CheY3*. Ni-NTA affinity purification using FLAG-tagged FliM (FliM-FLAG) and FLAG-tagged FliN (FliN-FLAG) to pull down HisCheY3 or HisCheY3* (CheY3^D79A^), respectively. HisCheY3 was co-purified with FliM-FLAG (**a**), but not with FliN-FLAG (**b**), suggesting that CheY3 does not bind on FliN. In contrast, more HisCheY3 protein was co-purified with FliM-FLAG with acetyl phosphate (**a**), and HisCheY3* was not co-purified with FliM-FLAG (**a**) or FliN-FLAG (**b**). These results indicate that CheY3 binds to FliM protein in a phosphorylation-dependent manner.

**Extended Data Fig. 7.**
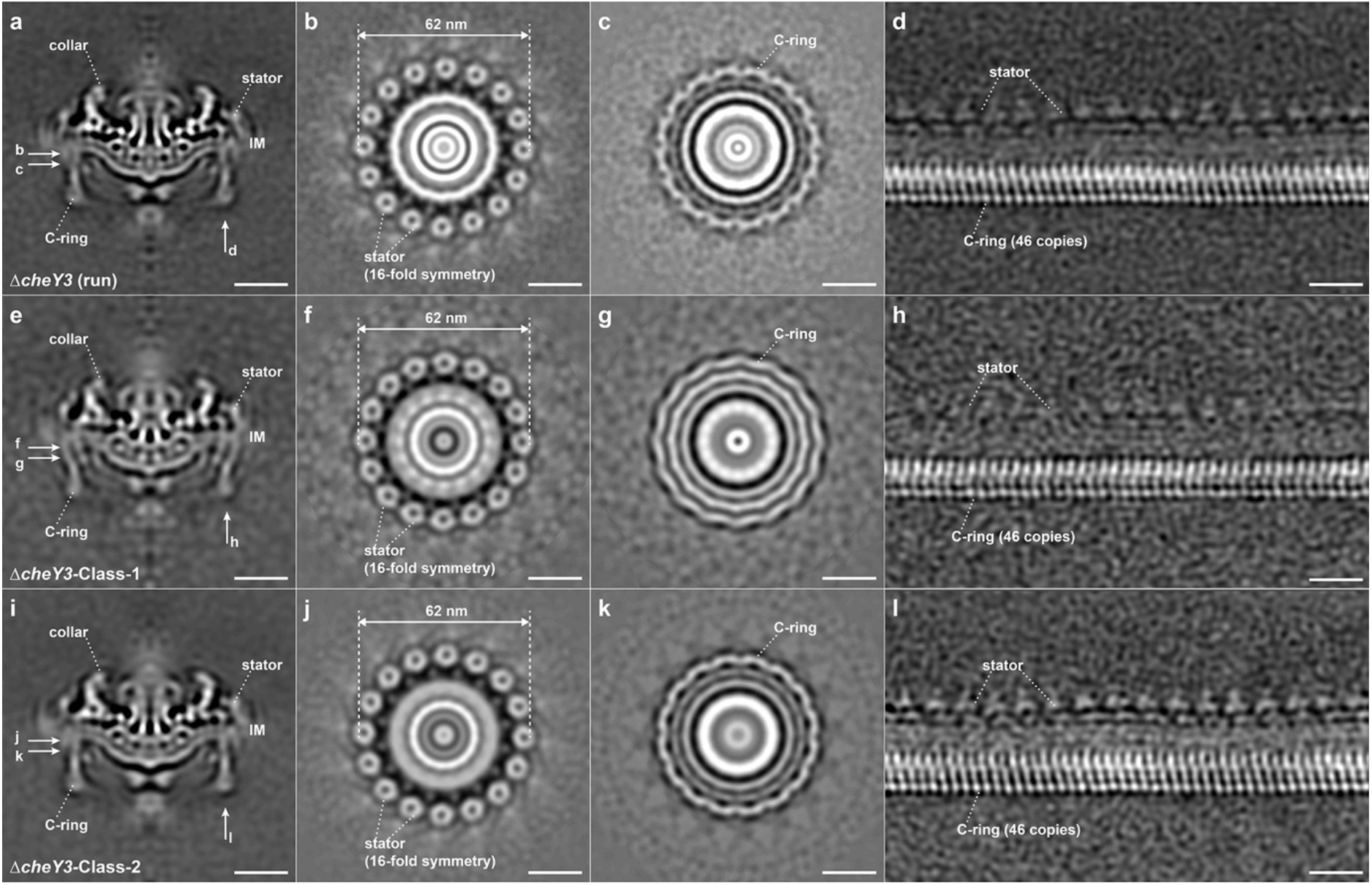
Motor structures in constantly running Δ*cheY3* cells. (**a**) A medial cross-section of an averaged structure in Δ*cheY3 B. burgdorferi* cell. (**b, c**) Cross-sections show the stator ring and the C-ring, respectively. (**d**) Focused structure of the C-ring (unrolled along the central rod). Two distinct classes in the Δ*cheY3* cells are named as Δ*cheY3*-Class-1 (**e-h**) and Δ*cheY3*-Class-2 (**i-l**). Class-1 and Class-2 account ~45% and ~55% of all the Δ*cheY3* motors we used for current work, respectively. The stator structures in Class-1 and Class-2 (**f** and **j**) are quite similar, while the C-ring subunits (compare h with l) are tilted in different directions.

**Extended Data Fig. 8.**
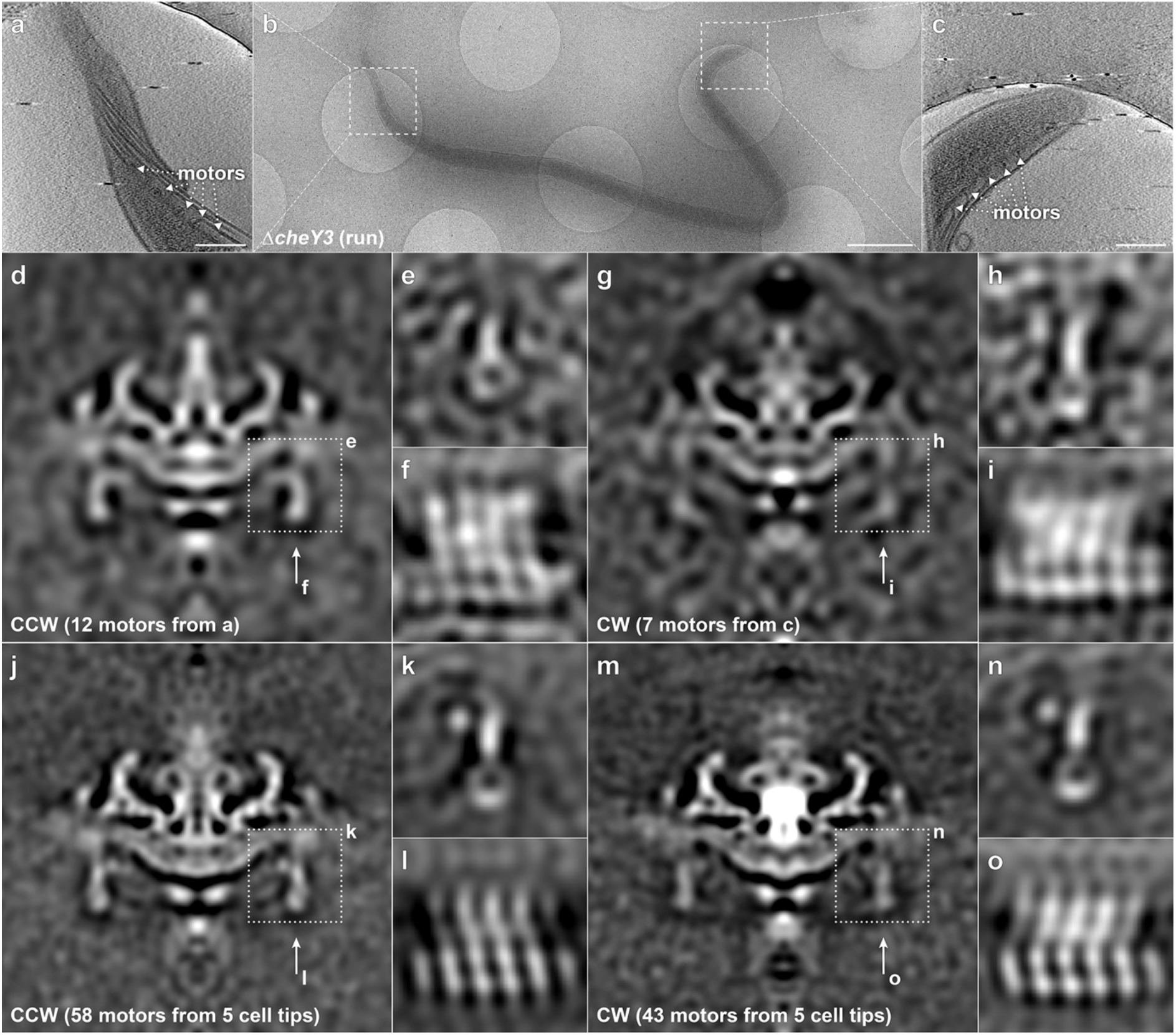
Motors adopt distinct conformations at the two cell poles in the same Δ*cheY3* cell. (**a**) A tomographic section from one cell tip showed in panel **b**. (**b**) An overview of one intact Δ*cheY3* cell. (**c**) A tomographic section from another tip of the same cell in panel **b**. The motors at each cell tip were aligned separately, then focused refined to the C-ring. (**d-f**) The motors from one cell tip have CCW conformation. (**g-t**) The motors from another tip appear to adopt CW conformation. (**j-l**) Averaged structure from motors located at one tip of five cells shows a better structure with CCW conformation. (**m-o**) Averaged structure from motors located at another tip of five cells shows a better structure with CW conformation. Bar = 200 nm in (**a, c**). Bar = 1 μm in (**b**).

**Extended Data Fig. 9.**
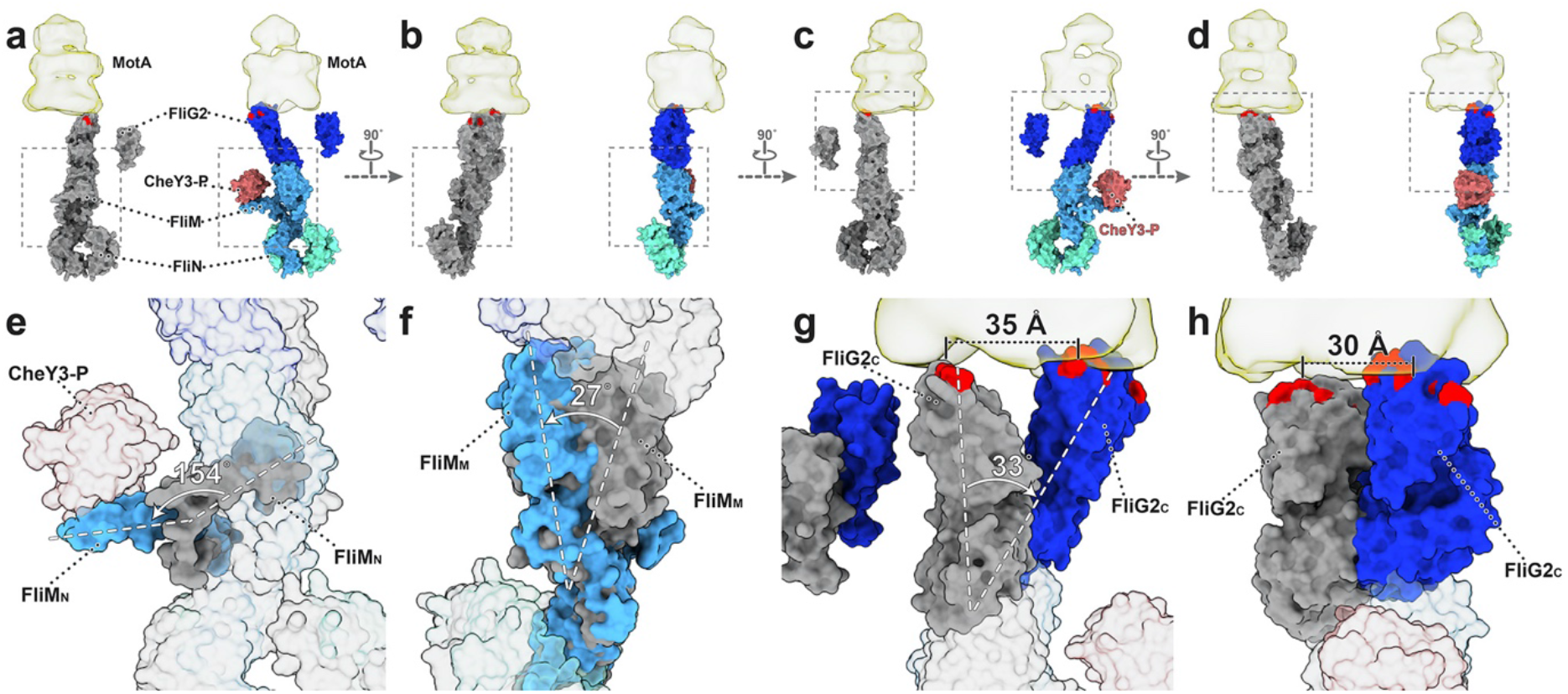
CheY3-P binding triggers conformational change. (**a-d**) Comparison between the C-ring models before (grey, top left in each panel) and after (colored, top right in each panel) CheY3-P binding. (**e**) The dash framed regions in panel **a** are overlapped to show their differences. The N-terminal domain of FliM (FliM_N_) folds out ~154° to interact with CheY3-P. (**f**) Binding of CheY3-P induces ~27° tilt of the FliM middle domain (FliM_M_). (**g, h**) FliG2 undergoes a large tilt and alters the interactions between FliG2 and MotA. The charged residues (Lys275, Arg292, Glu299, and Asp300) in the C-terminal domain of FliG2 (FliG2_C_) are colored in red.

**Extended Data Fig. 10.**
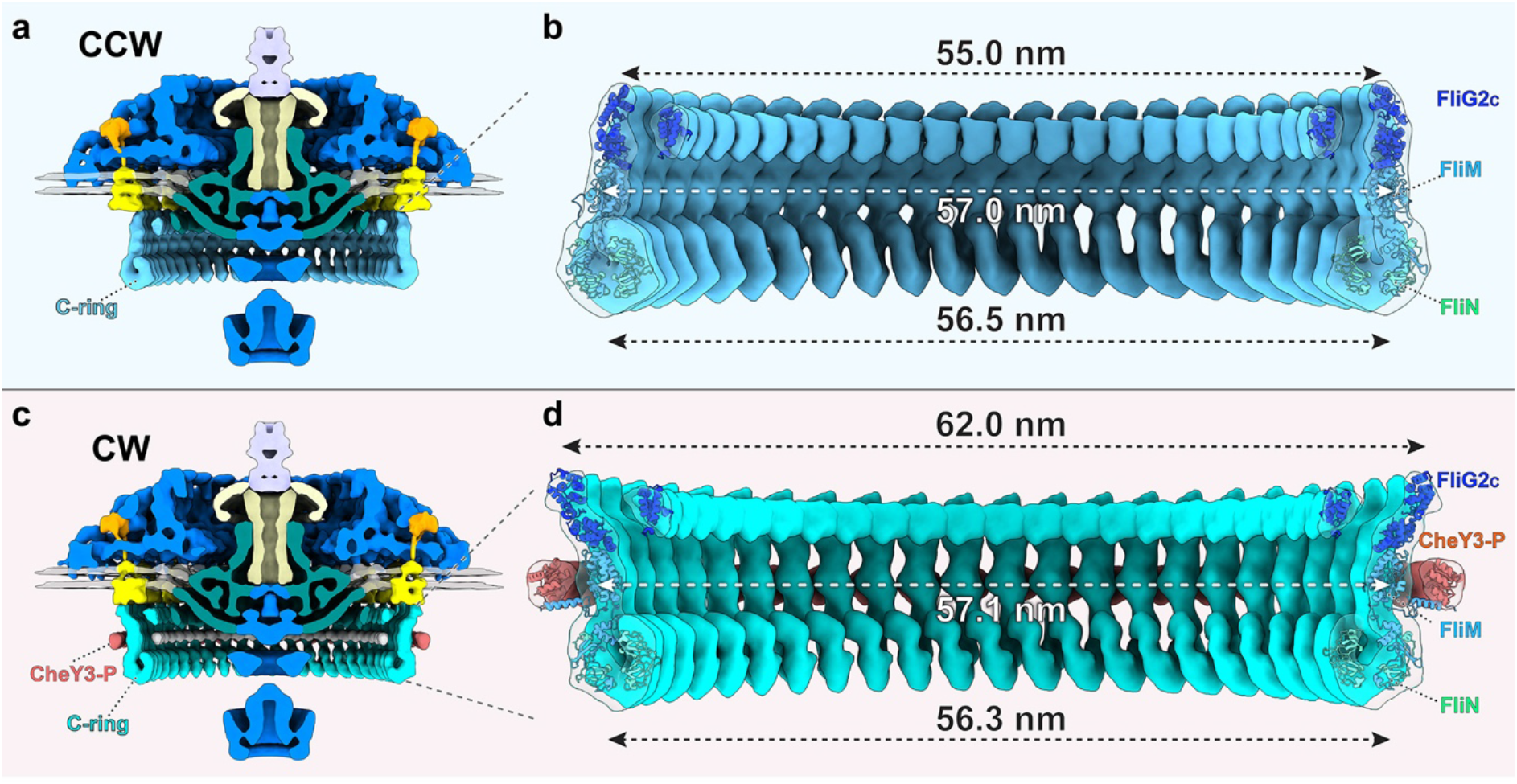
Comparison of the C-ring structures in CCW and CW rotation. (**a, b**) Diameters of the FliG2_C_, FliM and FliN rings in the C-ring with CCW rotation (Δ*cheY3*-class-2). (**c-d**) Diameters of the FliG2, FliM and FliN rings in the C-ring with CW rotation (Δ*cheX*).

**Extended Data Fig. 11.**
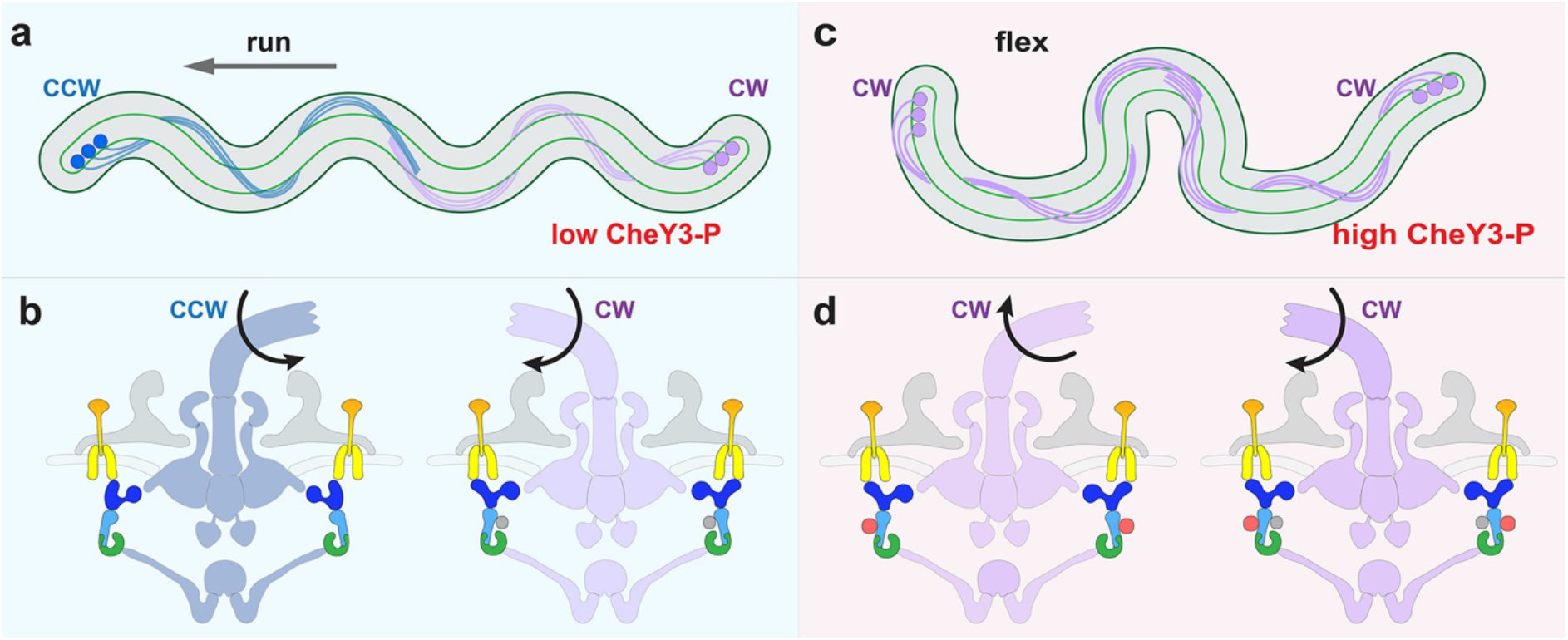
Motility model for *B. burgdorferi*. (**a, b**) In the default state, the concentration of CheY3-P is low, and the cell runs. The motors at the anterior cell pole rotate CCW, and the motors at the posterior cell pole rotate CW. Binding of unidentified proteins (grey circles at the inner side of the C-ring) to the C-ring at the posterior cell pole likely changes the motor to a CW conformation. (**c, d**) At high concentrations of CheY3-P, the CCW rotating motors switch to CW rotation, while the CW rotating motors keep turning CW. Thus, the motors at both cell poles rotate CW and the cell flexes. After the flex, the direction of flagellar rotation at the two poles can switch so that the cell reverses the direction of its run.

**Extended Data Table 1.**
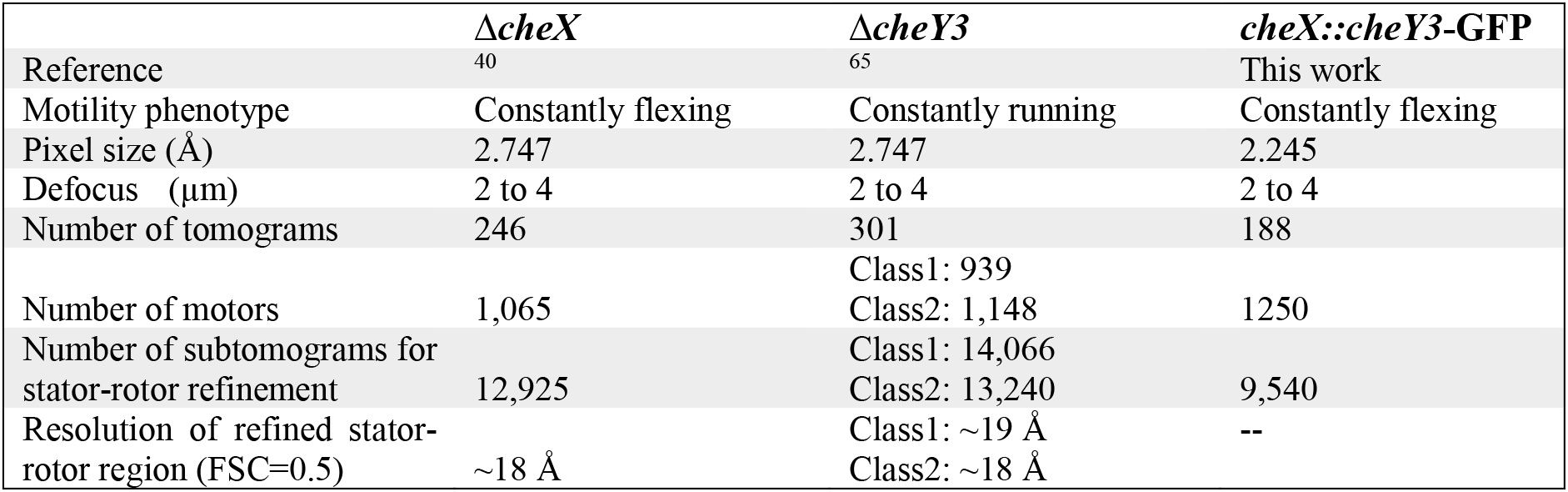
Strains and cryoET data used for this study.

**Extended Data Table 2.**
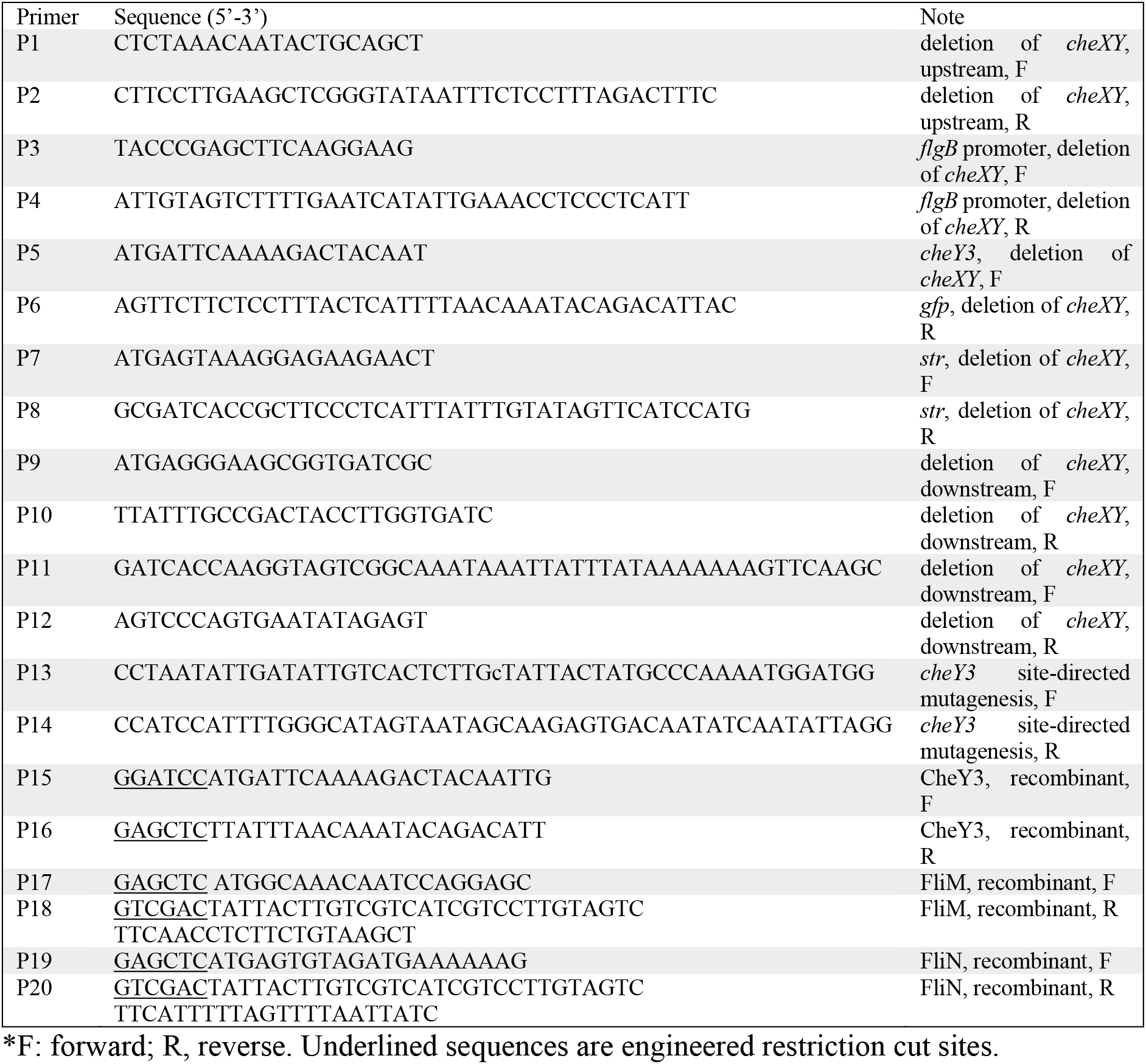
Oligonucleotide primers used in this study.

**Supplementary Video 1. Δ*cheX* cells flex in the video.**

**Supplementary Video 2. A refined structure of the motor in the Δ*cheX* mutant.**

**Supplementary Video 3. Δ*cheY3* cells constantly run in the video.**

**Supplementary Video 4. A class average of the motor in the Δ*cheY3* mutant.**

**Supplementary Video 5. Another class average of the motor in the Δ*cheY3* mutant.**

**Supplementary Video 5. Animation showing flagellar rotational switching in the Lyme disease spirochete.**

